# Sequencing-based quantitative mapping of the cellular small RNA landscape

**DOI:** 10.1101/841130

**Authors:** Jennifer F. Hu, Daniel Yim, Sabrina M. Huber, Jo Marie Bacusmo, Duanduan Ma, Michael S. DeMott, Stuart S. Levine, Valerie de Crécy-Lagard, Peter C. Dedon, Bo Cao

## Abstract

Current next-generation RNA sequencing methods cannot provide accurate quantification of the population of small RNAs within a sample due to strong sequence-dependent biases in capture, ligation, and amplification during library preparation. We report the development of an RNA sequencing method – AQRNA-seq – that minimizes biases and enables absolute quantification of all small RNA species in a sample mixture. Validation of AQRNA-seq library preparation and data mining algorithms using a 963-member microRNA reference library, RNA oligonucleotide standards of varying lengths, and northern blots demonstrated a direct, linear correlation between sequencing read count and RNA abundance. Application of AQRNA-seq to bacterial tRNA pools, a traditionally hard-to-sequence class of RNAs, revealed 80-fold variation in tRNA isoacceptor copy numbers, patterns of site-specific tRNA fragmentation caused by stress, and quantitative maps of ribonucleoside modifications, all in a single AQRNA-seq experiment. AQRNA-seq thus provides a means to quantitatively map the small RNA landscape in cells and tissues.

---

Next generation RNA sequencing platforms (RNA-seq) have advanced functional genomics by facilitating discoveries of new RNA species, gene annotations, and quantitative analysis of gene expression.^1, 2^ RNA-seq has played a central role in unravelling the complexity of RNA function by facilitating quantitative profiling of the transcriptome as a function of cell state. Although a variety of RNA-seq methods provide precise and accurate analysis of changes in transcript abundance *<u>between</u>* samples or conditions, their major drawback is the inability to accurately quantify small RNA species *<u>within</u>* a sample. This is partly rooted in biased ligation of sequencing linkers to the 3’- and 5’-ends of RNAs, which varies by 10^3^-fold depending upon on the identity of the terminal nucleotides^3-7^ and causes 10^6^-fold artifacts in sequencing read counts.^5, 8, 9^ Highly structured and heavily modified RNA molecules, such as tRNAs, further challenge the quantitative accuracy of RNA-seq.^10, 11^ These structural features cause polymerase fall-off during cDNA synthesis, which prevents detection of the transcripts.

A variety of specialized RNA-seq methods, many limited specifically to either miRNA or tRNA only,^12-14^ attempt to minimize ligation bias using linkers with randomized terminal nucleotides and molecular crowding agents to enhance ligation^4^ and to reduce polymerase fall-off during reverse transcription (RT) with two-step ligation^8^ and removal of some methyl modifications with AlkB.^15-17^ Although polymerase fall-off effects are minimized, residual ligation biases persist and lead to “jackpot” sequences.^8^ For example, the template-switching reverse transcriptase TGIRT used in DM-tRNA-seq^16^ is biased by the identity of the overhanging nucleotide in the adapter strand.^18^

The problem with existing RNA-seq techniques is that none have been systematically engineered to optimize ligation and amplification efficiencies or validated for quantitative accuracy and lack of bias artifacts. Furthermore, none are broadly applicable to all RNA species. Here we describe AQRNA-seq, an RNA-seq method that enables absolute quantification of all RNA species in a sample by providing a direct, linear correlation between sequencing read count and RNA abundance. The library preparation and data mining algorithms have been rigorously validated by multiple orthogonal approaches. Application of AQRNA-seq to the dormancy response of *Mycobacterium bovis* BCG revealed large variations in tRNA copy numbers, tRNA fragmentation, and tRNA modification location and abundance within and among samples along a time course of stress-induced mycobacterial persistence.

## Results

### AQRNA-seq design and optimization

The AQRNA-seq workflow (**Figure 1a**) maximizes ligation capture of RNAs using novel adapters (linkers) and minimizes RT fall-off with two-step linker ligation and AlkB treatment. Adapter ligation (“linker 1”) begins at the 3’-end, with two randomized nucleotides incorporated at the 5’ end of linker 1 to maximize ligation efficiency based on known strong biases in T4 ligase activity.^19^ Linker 1 is also a DNA oligo to facilitate removal of unligated linker with RecJ, a single-stranded-DNA-specific 5’→3’ exonuclease, which leaves the hybrid RNA-DNA product intact.^20^ A 50:1 excess of linker 1 resulted in >90% ligation efficiency (**Fig. 1b**).

**Figure 1.**
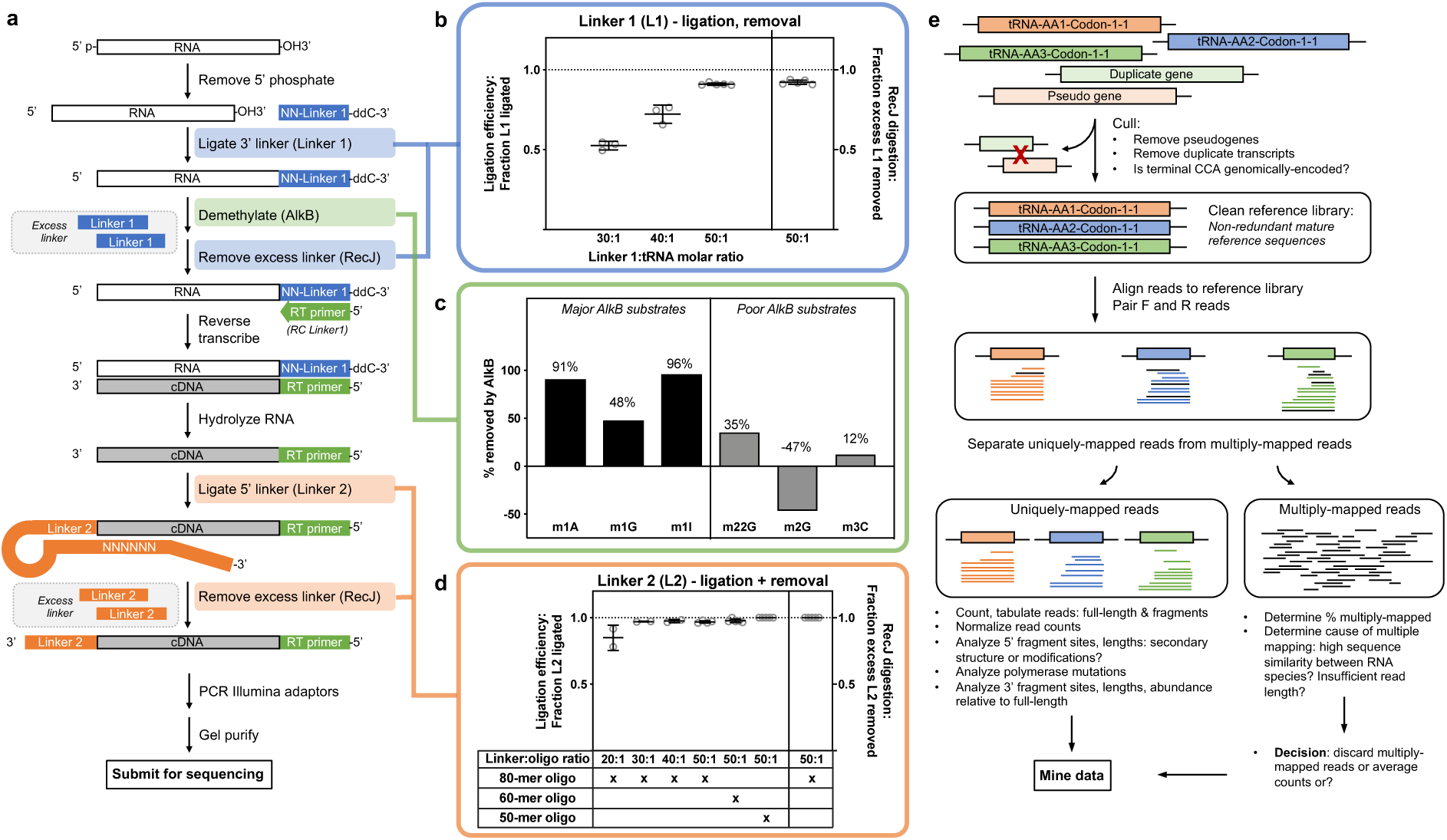
Overview of AQRNA-seq. (**a**) Library preparation workflow. (**b**) Control/optimization experiments for steps in the workflow. Top panel: Linker 1 ligation and removal proceeds with greater than 90% efficiency at a linker to tRNA molar ratio of 50:1. Middle panel: LCMS measurements of AlkB demethylation efficiencies for various RNA modifications. Bottom panel: Linker 2 ligation proceeds with near stoichiometric efficiency at linker to tRNA molar ratios of 30:1 or above. A 50:1 ratio is used in the method. Linker 2 removal is 100% efficient under those conditions. (**c**) Computational analysis workflow. Reads are mapped against a non-redundant reference genome. cDNA libraries generated and sequenced according to the AQRNA-seq, a paired-end protocol, will yield two FASTQ files per library – one containing the forward reads, and one containing the reverse reads. After alignment, multiply-mapped reads are mined separated from uniquely-mapped reads for abundance and coverage information.

**Figure 2.**
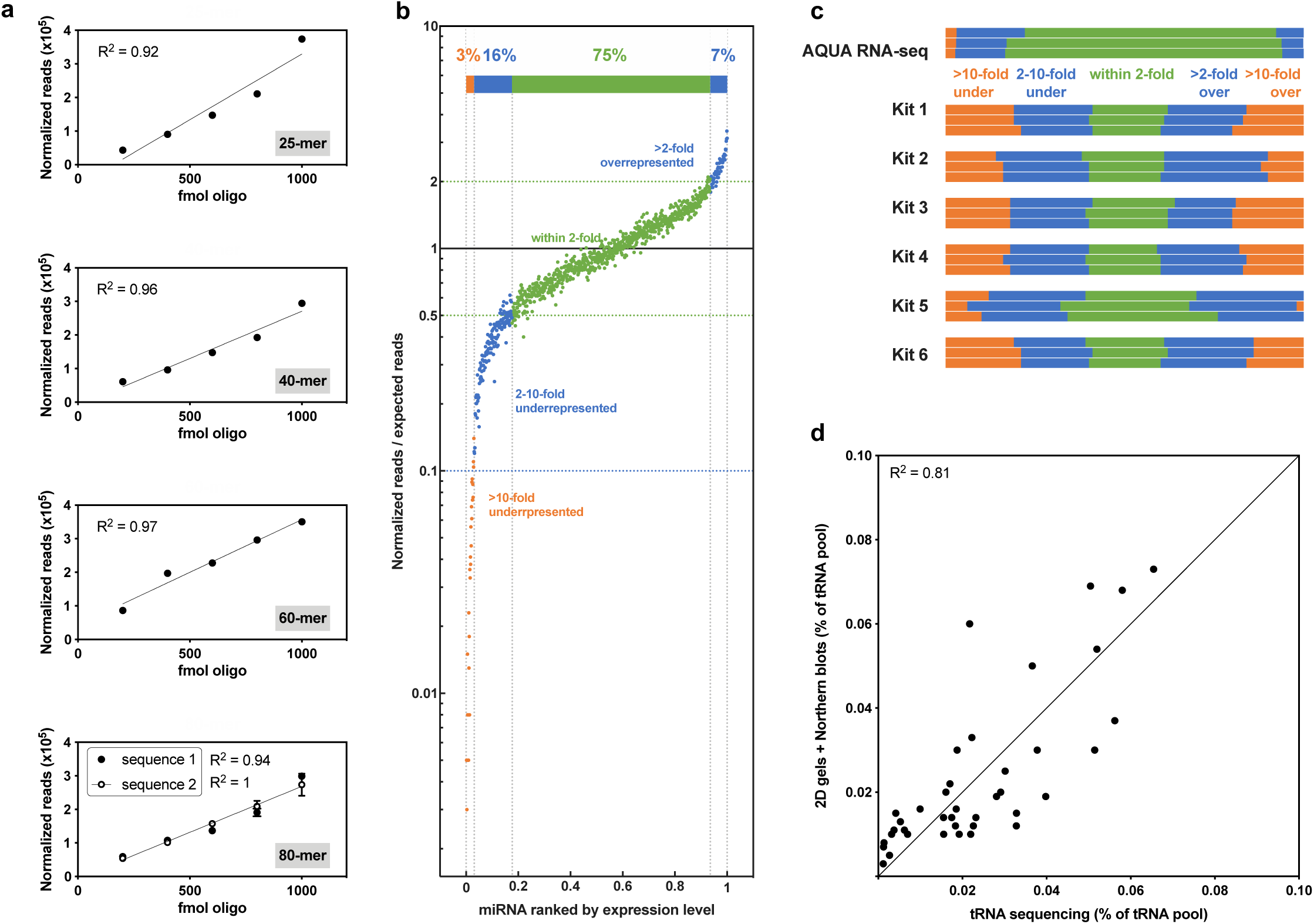
Validation of AQRNA-seq. (**a**) Oligo spike-in studies. Five RNA oligos between 25 and 80 nucleotides in length were prepared for sequencing at different concentrations. The sequencing response per fmol of oligo is linear with amount of material present. (**b**) An equimolar mixture consisting of 963 synthetic microRNAs was sequenced using AQRNA-seq. Sequencing reads for each microRNA were normalized to their expected values and sorted. The colored bars and dotted lines indicate the percentage of the total mixture came within 2-fold and 10-fold of their expected abundance. (**c**) A comparison of microRNA libraries prepared using the AQRNA-seq protocol and various commercially-available small RNA sequencing kits shows significantly better performance for AQRNA-seq in terms of quantitative accuracy. Data for Illumina TruSeq Small RNA Library Prep Kit, Lexogen Small RNA-seq Library Prep Kit, NEBNext Small RNA Library Prep Set for Illumina, BioO POWERFLEX, QIAseq miRNA Library Kit, and Trilink CleanTag Small RNA Library Prep Kit taken from Herbert et al.^47^ (**d**) Correlation of tRNA quantification results using AQRNA-seq versus data derived from 2D gel electrophoresis and northern blotting by Dong et al.^26^

The ligated RNA can then be treated with AlkB to reduce levels of RT-blocking methyl modifications.^15^ Though this step is not essential for capturing all RNA sequences in a sample, it can provide information about the identities of polymerase-interfering modifications. Optimized buffer conditions and AlkB concentration allowed removal of m^1^A (90%), m^1^G (48%), and m^1^I (96%). Only 12% of m^3^C was removed, which is contrary to previous observations^16^, and m^2,2^G was reduced by only 35%, apparently demethylated to AlkB-resistant m^2^G (**Fig. 1c**).^15^ After demethylation, unligated linker 1 is removed with RecJ. Optimized conditions resulted in >99% linker removal (**Fig. 1b**), obviating the need for HPLC purification of the RNA-DNA product.

RT is now accomplished with a DNA primer complementary to the 3’ DNA linker 1 and the resulting cDNA is ligated at its 3’-end to a custom DNA adapter (linker 2) using T4 DNA ligase. Linker 2 possesses a hairpin, a random hexamer sequence, and a downstream primer binding site for subsequent amplification. The random hexamer in linker 2 acts as a splint to enhance ligation with the cDNA (**Fig. 1d**), which was optimized to nearly 100% with a 50:1 linker excess (**Fig. 1d**). Excess linker 2 is removed with RecJ and PCR amplification was performed with primers that complement linker 2 and incorporate standard Illumina anchor and barcode sequences for subsequent sequencing.

### AQRNA-seq data processing workflow

Rigorous processing of the RNA sequencing data represents another step critical for quantitative accuracy. To this end, we developed a customized workflow for processing AQRNA-seq sequencing data, illustrating the main points here for bacterial tRNA sequences (**Fig. 1e**). Similar customization is necessary for other RNA species and for tRNAs from higher eukaryotes. Building on several commonly used software packages for traditional RNA-seq analyses (see Methods), workflow customization begins with curation of the reference sequences used for aligning sequencing reads. Here we focus on bacterial tRNAs, with considerations applying to all transcripts in any organism. Reference sequences are first culled of duplicate genes and pseudogenes that lead to ambiguous assignment of reads. Similar consideration must be given to post-transcriptional processing, such as trimming and processing of 5’- and 3’-termini of primary tRNA transcripts as well as tRNA modifications.^21^ For mycobacteria, the 3’-CCA of each tRNA is variably genomically encoded or added post-transcriptionally.^22^

The resulting non-redundant reference library is then used to align forward and reverse sequencing reads, first separating uniquely aligned reads from reads matching multiple sequences in the library. Even with careful library curation, high sequence similarity or insufficient read length can result in multiply-mapped reads. We arbitrarily set a 10 nt read length filter to maximize alignments. For closely-related reference sequences, multiply-mapped reads can be resolved by collapsing read assignments to broader “families” (**Fig. 1e**), with subsequent determination of the proportion of multiply-mapped reads and the cause of multiple mapping. These considerations rationalize a decision to discard, average, or sum the read counts from multiple, closely-related reference sequences.

Finally, the read count for full-length sequences and fragments is tabulated from the curated set of mapped reads. Data are normalized either to the total number of reads in each barcoded sample or to an internal RNA standard to account for sample-to-sample variation in input RNA as well variation in sample pooling prior to sequencing. For tRNAs, the resulting dataset can then be visualized using a graphical presentation that reveals abundance, fragmentation sites, and modification sites in a single view, as illustrated shortly for stress-induced bacterial persistence (**Figs. 3-7**).

**Figure 3.**
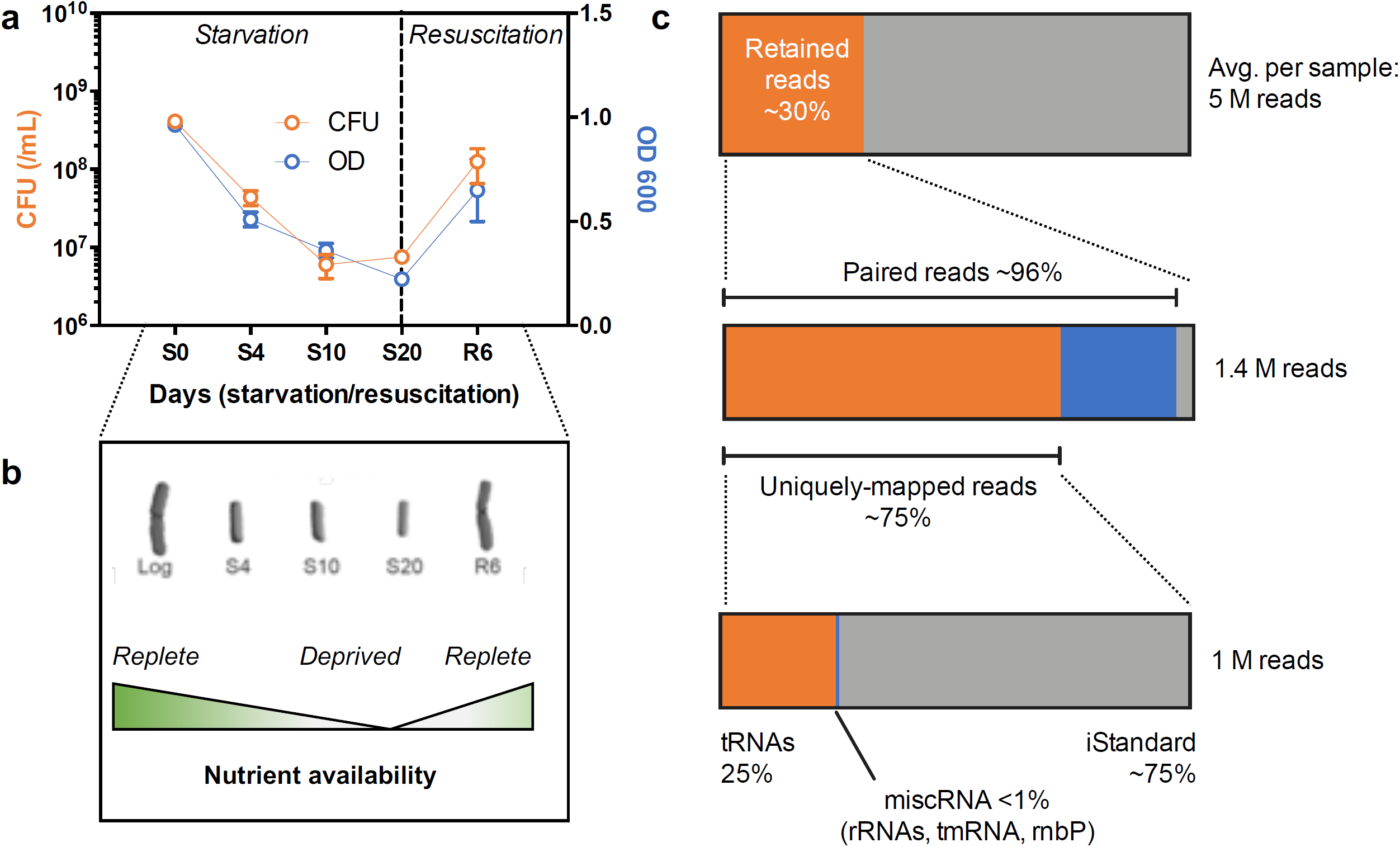
Application of AQRNA-seq to quantify tRNA pool dynamics during stress-induced persistence in *M. bovis* BCG. (**a**) Growth curves of showing approximately two logs of cell killing from day 0 to day 20 of complete nutrient deprivation followed by regrowth after resuscitation. (**b**) *In vitro* model of nutrient deprivation in *M. bovis* BCG. (**c**) A breakdown of the distribution of reads after size-selection, pairing of forward and reverse reads, and mapping. Of uniquely mapped and paired reads, 75% mapped to the internal standard sequence while 25% mapped to mature tRNA sequences.

### Method validation shows a linear relationship between read count and RNA abundance

Three different approaches were used to test the precision and accuracy of AQRNA-seq. First, five synthetic RNA oligos with varying lengths (25-80 nt) were mixed at varying molar ratios and used as input RNA for library preparation. After sequencing, we found that read counts for each oligo varied directly and linearly with input RNA abundance (r^2^ 0.92-1; **Fig. 2a**), with an averaged sequencing response (slope) of ∼300 ± 50 reads per fmol of input RNA. This demonstrates minimal sequencing bias for quantity or length of input RNA.

To test for sequence-dependent ligase or polymerase biases, we used AQRNA-seq to prepare libraries from the Miltenyi miRXplore Universal Reference consisting of 963 equimolar miRNA sequences (sourced from miRBase^23^), which ranged in length (16-28 nt) and possessed all 16 possible dinucleotide combinations at 3’ and 5’ termini. Resulting sequencing reads for each miRNA were counted and divided by total counts for all detected miRNAs to obtain a “normalized read count”. Total counts were also divided by 963 (the number of detected miRNAs) to obtain the “expected read count”, assuming all species are equimolar. A read ratio was then calculated by dividing “normalized” by “expected”. A plot of all 963 read ratios ranked from lowest to highest showed that ∼75% fell within two-fold variation of expected abundance (**Fig. 2b**). The number of jackpot and dropout miRNAs (those with normalized read ratios >10-fold higher or lower than expected) was <3% of the total miRNA mixture. No other library preparation method resulted in greater than 40-50% of miRNAs falling within the two-fold threshold.^24^ Based on this comparison and additional reports using the miRXplore reference,^5, 18, 25^ AQRNA-seq is the most quantitatively accurate RNA sequencing workflow currently available.

The third validation study tested AQRNA-seq performance against the analysis of the *E. coli* K12 tRNA pool by two-dimensional gel electrophoresis and Northern blotting by Dong et al.^26^ Applying AQRNA-seq to small RNAs (<200 nt) from *E. coli* K12 strain BW25113 (derived from W1485 used by Dong et al.^26^), the total expressed levels of 45 tRNAs (summed read counts for full-length and truncated sequences) were compared to the 46 tRNAs identified by Dong et al.^26^ Excluding one outlier, there was strong agreement (r^2^ = 0.81) between the two orthogonal approaches (**Fig. 2d**).

### tRNA landscape dynamics in stress-induced bacterial persistence

AQRNA-seq was now applied to quantify the dynamics of a particularly challenging set of targets: the 45 tRNAs in the *Mycobacterium bovis* BCG model for stress-induced, non-replicative, antibiotic-resistant state of persistence in tuberculosis.^27-29^ Total small RNA (<200 nt) was isolated along the time course of BCG persistence caused by complete nutrient deprivation (**Fig. 3a,b**), with ∼1% of the bacterial population surviving as persisters after 20 days in PBS and restoration of growth with transfer to nutrient-rich medium (**Fig. 3a**). Processing of the AQRNA-seq read data for one BCG starvation study illustrates the workflow points made earlier. In this highly-multiplexed sequencing experiment involving triplicate samples from each of 5 time points during starvation and resuscitation, an average of 5 million raw sequencing reads was obtained for each sample (**Fig. 3c**). After size-selection and adapter trimming, the majority (75%) of the remaining reads consisted of uniquely-mapped, paired reads when mapped to the full set of mature BCG RNA sequences (**Fig. 3c**). Of these, another 75% mapped to an 80-nt internal standard that was added in large excess in this experiment, while remaining reads mapped to reference library tRNA sequences (**Fig. 3c**). A much lower level of internal standard allows detection of rare RNA species, such as tRNA fragments illustrated in **Figure 4e** as discussed shortly. To account for variation introduced by input RNA and sample processing, reads originating from a single sample can be normalized to a spiked-in standard (80 nt here). Comparison between samples is facilitated by expressing RNA read counts as either a percentage of total aligned reads or total aligned tRNA reads within each sample. For example, the abundance of a single isoacceptor can be expressed as a proportion of the complete isoacceptor pool in a sample and this value can be tracked and compared across samples derived from various treatment conditions. This normalization process applies to any RNA species of interest.

**Figure 4.**
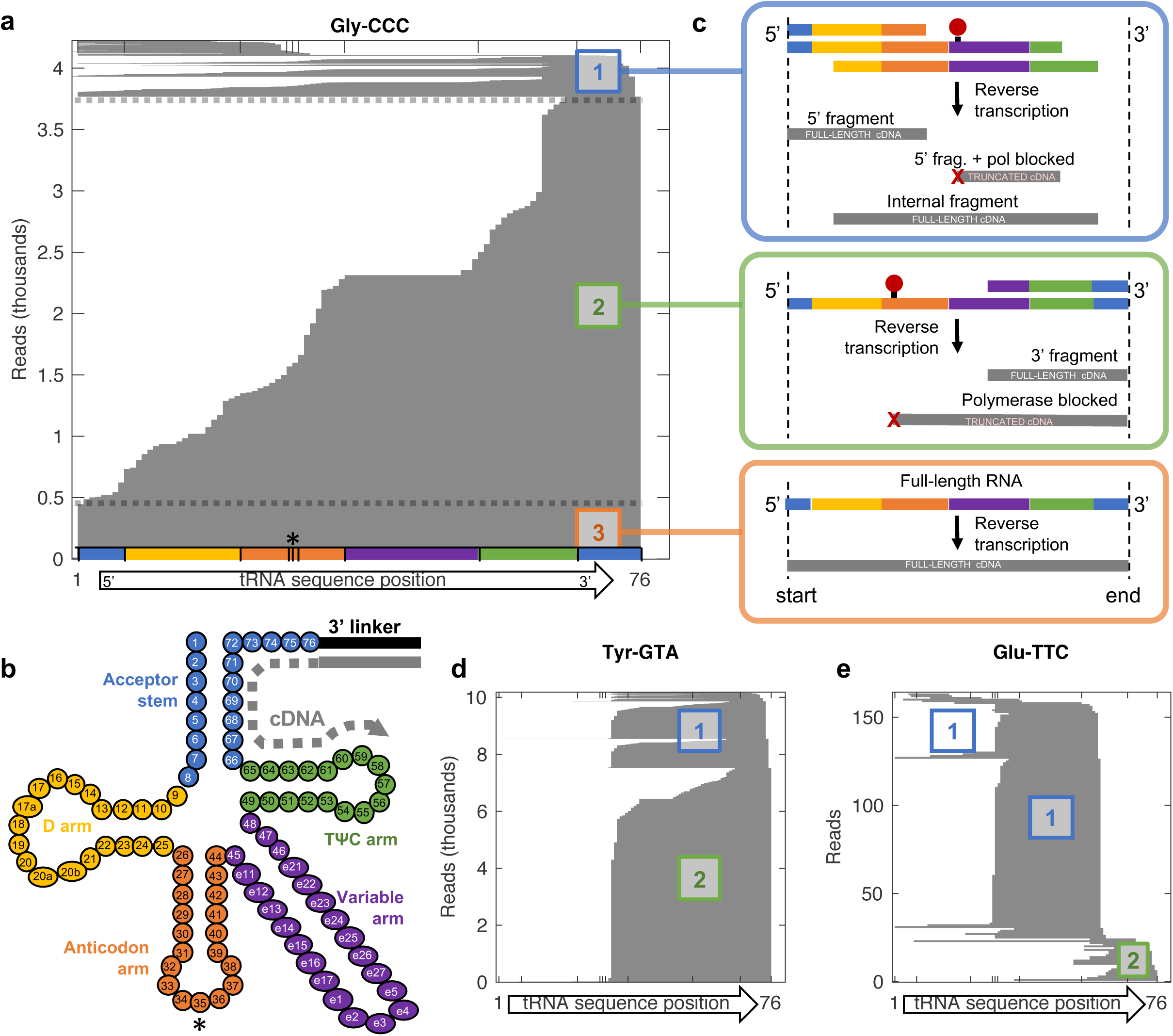
AQRNA-seq quantifies the complete landscape of full-length and fragmented tRNAs, with single-nucleotide resolution of cleavage sites and modifications. (a) Sequence alignment plots reveal the abundance of quantitative information provided by AQRNA-seq. This is illustrated for tRNA Gly-CCC, with a stacked horizontal bar graph of the sequence alignment showing the start and end position of reads aligning to tRNA Gly-CCC. The numbering of tRNA nucleotides follows the standard nucleotide number system with positions 1 and 76 reflecting the 5’ and 3’ termini respectively. The anticodon is indicated by three vertical lines and an asterisk. The complete set of alignment plots for the 45 tRNA isoacceptors in *M. bovis* BCG is shown in **Supplementary Figure S1**. (**b**) Schematic showing site of linker 1 attachment and direction of reverse transcription along the tRNA sequence annotated using the Sprinzl coordinate system.^30^ (**c**) The aligned reads fall into three different categories which have different interpretations – see text. (**d**) Example alignment plot for tRNA Tyr-GTA showing polymerase blockage downstream of the anticodon resulting in a lack of type 3 (full-length) reads. (**e**) Example of an alignment plot to tRNA Glu-TTC showing polymerase blockage close to the anticodon and an enrichment of fragments aligning inside the 3’-end of the tRNA, resulting in an increased number of type 1 reads.

The resulting BCG dataset was mined for information about starvation-induced changes in tRNA expression, tRNA fragmentation patterns, and the locations of modified nucleosides in individual tRNAs. These tRNA features are best visualized graphically in horizontally stacked alignment plots (**Fig. 4a**), in which the start and end positions of each sequencing read are aligned along an X-axis annotated using the Sprinzl coordinate system for tRNA structure (**Fig. 4b**).^30^ This plot format readily informs about the state of each tRNA isoacceptor, with the 3’ end of a read reflecting where the source RNA species was captured by linker 1 (**Fig. 4c**). The complete set of alignment plots for 45 unique, annotated tRNA species in BCG is shown in **Figure S.1.** As a first observation, the height off the stack is directly proportional to the total number of expressed transcripts. The graph can then be split into three sections. The bottommost portion consists of full-length reads – reads that span the entire tRNA sequence (**Fig. 4b**). These “type 3” reads originated from full-length tRNA molecules. The top section contains reads that do not reach the 3’ end of the tRNA (“type 1” reads; **Fig. 4b**) and correspond to 5’ tRNA fragments, with the 3’ linker ligated to the 5’ end of the break site (**Fig. 4c**). The middle section contains reads that start at the 3’ end of the tRNA but do not reach the 5’ end (“type 2” reads; **Fig. 4b**). These reads correspond to tRNA molecules that were fragmented to leave behind a 3’ portion, or to full-length tRNAs for which cDNA synthesis was prematurely truncated due to polymerase fall-off. Analysis of 3 tRNA isoacceptors by northern blotting using 5’ and 3’ probes (**Fig. S2**) suggests that the majority of the type 2 reads are in fact full-length tRNAs as the vast majority of bands detected using the 3’ probe were full-length. This is consistent with the relatively low level of 5’ tRNA fragments in stack plots for all of expressed tRNA molecules in BCG (**Fig. S1**).

### Starvation-induced persistence remodels the tRNA landscape in BCG

AQRNA-seq provides a global view of changes in the RNA landscape caused by stress. In starved BCG, the abundance of individual tRNAs – here defined as the sum of all reads that aligned to a particular tRNA species – spanned a large range. In most samples, tRNA Lys-CTT and tRNA fMet-CAT were the most highly expressed tRNAs, together totaling ∼20% of the tRNA pool during log growth, whereas tRNA Ser-GGA was ∼80-times lower, comprising only 0.1-0.3% of the tRNA pool (**Figs. 5a, S1**). In the transition from rich medium to starvation over 20 days, the abundances of several tRNAs are significantly altered (**Fig. 5a**,**b; Fig. S1**). For example, tRNA Ser-CGA and tRNA His-GTG drop significantly in early starvation and rise again during late starvation and resuscitation. tRNA Leu-CAG and tRNA Thr-GGT show the opposite pattern, increasing during starvation and returning to S0 levels during resuscitation.

**Figure 5.**
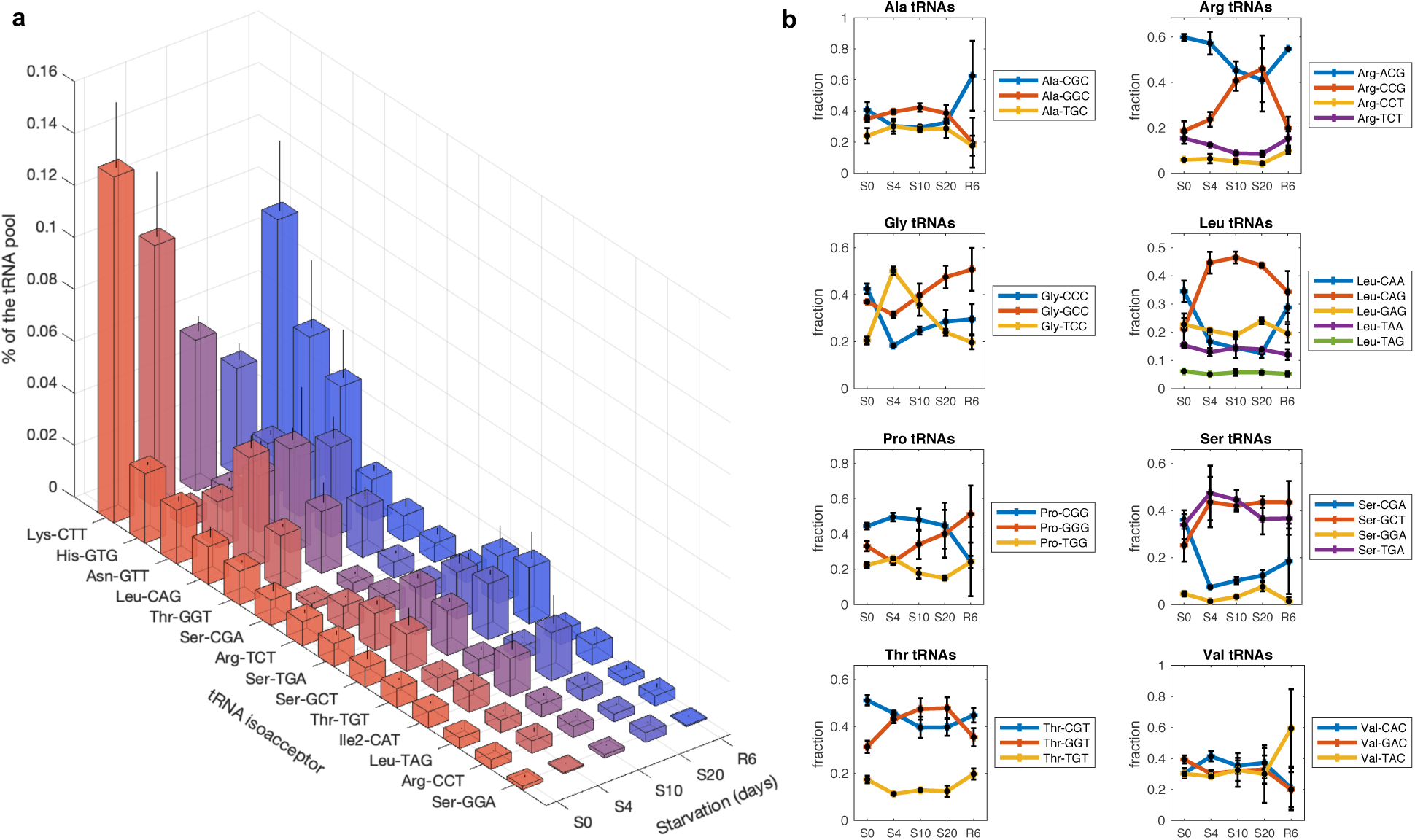
AQRNA-seq quantifies the tRNA pool dynamics during stress-induced persistence in *M. bovis* BCG. (**a**) Plots showing the normalized abundance of all expressed BCG tRNAs throughout the time course of starvation. The green line represents the mean of tRNA abundances at the S0 timepoint. (**b**) Plots showing abundances of individual isoacceptors (all reads aligning at the 3’ relative to the total set of tRNAs that carry the same amino acid.

Starvation also induced significant shifts among the isoacceptors associated with a specific amino acid (**Fig. 5b**). In BCG, there are 7 amino acids for which there is a single isoacceptor (Trp, Asp, Asn, Cys, His, Phe, Tyr), 5 have 2 isoacceptors, and 8 have ≥3 tRNA isoacceptors. Illustrating the latter, Ser, Thr, and Leu are specified by 6, 4, and 6 synonymous codons, respectively, and these codons are read by 4, 3, and 5 different tRNA species, respectively.^31^ **Figure 5b** shows starvation-induced changes in fractional abundance of isoacceptors for these amino acids. Prior to starvation (S0), tRNA Ser-CGA, tRNA Ser-TGA and tRNA Ser-GCT comprise 36%, 34%, and 25% of the Ser isoacceptor pool, respectively. At 4 days of starvation (S4), the abundance of tRNA Ser-CGA drops to 7% while tRNA Ser-TGA and tRNA Ser-GCT surge to 47% and 44%, respectively. For Thr, tRNA Thr-CGT and tRNA Thr-GGT represent 51% and 31%, respectively, during log growth (S0), but flip to 40% and 47% at S10, and 40% and 48% at S20, before returning to pre-starvation levels during resuscitation. These data illustrate the dynamics of individual tRNAs resulting from stress-induced changes in gene expression or tRNA degradation.

AQRNA-seq also captures the dynamics of tRNA fragmentation and degradation, as occurs in tRNA maturation and quality control,^32-34^ small RNA regulation of gene expression, and toxin-antitoxin systems.^35-40^ It is difficult to differentiate 5’-degradation of full-length tRNA from polymerase fall-off as both generate a fragment that aligns at the 3’ end of tRNA (“type 2” reads, **Fig. 4b**). However, 3’-degradation and endonucleolytic cleavage generate fragments with 3’ ends positioned inside the reference sequence (“type 1” reads, **Fig. 4b**).

An extreme case is illustrated by tRNA Glu-TTC. While ∼80% of the reads aligned with the 3’ end in log growth, ∼80% of the reads at S4 had 3’ ends located at position 58 in the TΨC arm of the tRNA (**Fig. 4b**). These tRNA fragments appeared as a gel band present in the S4 sample but much fainter in the log sample (**Fig. S3b**). The absence of a corresponding number of short (15-20 nt) tRNA fragments representing the 5’-side of the cleaved tRNA (type 2 reads, **Fig. 4e**) could result from degradation of the 3’-fragments. AQRNA-seq thus provides quantitative information about tRNA fragmentation in addition to total expressed tRNA copy numbers and, as discussed next, the locations of modified nucleosides and secondary structures in RNA.

### Quantitative mapping of RNA modifications and secondary structure

Nearly all forms of RNA contain post-transcriptional modifications, with over 120 structures known.^41^ tRNAs are particularly heavily modified to the extent of ∼10% of the component nucleotides, on average.^10^ In some cases, modifications interfere with the fidelity and processivity of the reverse transcriptase during RNA-seq library preparation, which can be exploited to map modification positions based the resulting sequence data. ^15, 42, 43^ AQRNA-seq detects RT defects as read pile ups at specific sites along the RNA sequence. We illustrate this behavior with small RNA isolated from log-growing *E. coli*, for which there is detailed information about the types and locations of tRNA modifications.^41^ As shown in **Figure 6a**, several tRNAs had high levels of polymerase stops at positions 38 and 48. By overlaying the sequencing stop positions on tRNA modification maps for *E. coli* in the Modomics database,^41^ we determined that these two positions are proximal to known tRNA modifications. One subset of 9 tRNAs had 31-83% of the mapped reads stop at position 48, which abuts the modification 3-(3-amino-3-carboxypropyl)uridine (acp^3^U) at position 47 (orange boxes on right in **Fig. 6a**). With U-*N*^3^ involved in base-pairing and *N*^3^-methyluridine (m^3^U) previously reported to block RT,^44^ it makes sense that acp^3^U would also prevent polymerase procession. tRNAs with NNA anticodons had 48-69% of the reads end at position 38, which is adjacent to *N*^6^-isopentenyladenosine (i^6^A) and its hypermodified derivatives (e.g., 2-methylthio-*N*^6^-isopentenyladenosine, ms^2^i^6^A) in the anticodon loop (yellow boxes on left in **Fig. 6a**). Modifications involving the *N*^6^ amine of adenosine are not known to block RT, but *N*^6^-methyladenosine induces pausing.^44^ It is also plausible that the bulky size of the isopentenyl modification combined with the sharp, three-dimensional turn of the anticodon hairpin interferes with polymerase activity.

**Figure 6.**
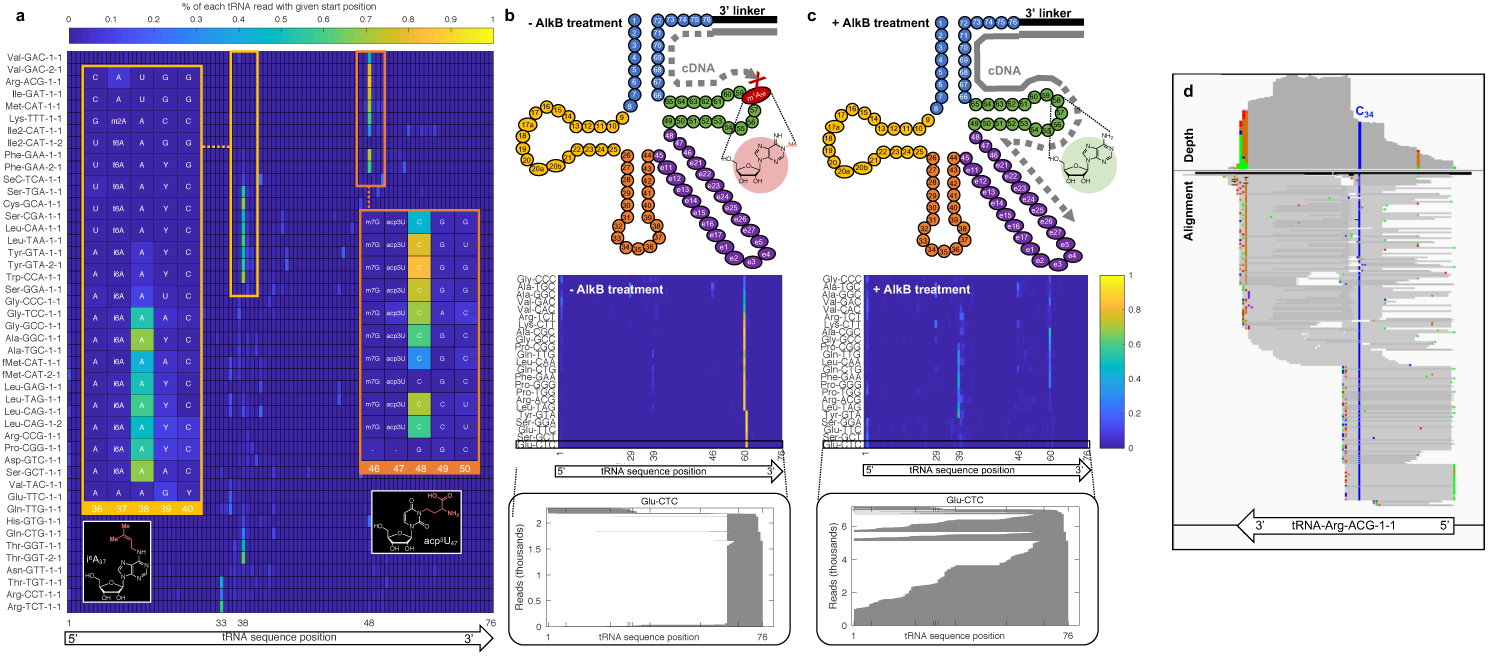
Application of AQRNA-seq for quantitative mapping of the tRNA epitranscriptome. Many RNA modifications block 3’-to-5’ reverse transcriptase-mediated cDNA synthesis or result in mutations in AQRNA-seq (gray line and arrow in panel **b**), which can be exploited to quantitatively map the modifications. (**a**) Subsets of *E. coli* tRNAs exhibit similar reverse transcriptase blockages at positions 38 and 48. The heat map shows the percentage of sequencing reads for which reverse transcription ended at a specific location in the tRNA sequence (columns) for each tRNA isoacceptor (rows). The two positions showing the most significant accumulations of polymerase blockade are noted with a small orange or red box. The RNA sequences surrounding these positions are shown in the larger boxes that magnify the sequence location, with specific modified nucleotides noted based on existing maps of *E. coli* tRNA modifications. In the orange boxes, the 8 tRNA species showing polymerase blockade at position 38 reveals all possess i^6^A at position 37. In the red boxes, the 10 tRNAs showing polymerase blockade at position 48 all possess acp^3^U at position 47. (**b**) In the absence of AlkB treatment, cDNA synthesis is blocked by m^1^A at position 58 in nearly half of all BCG tRNAs, which is reflected by the high proportion of aligned reads that do not extend past position 58 in the heatmap of read start positions (light blue to orange line in heat map similar to panel **a**). This is illustrated for tRNA Glu-CTC in the gray stack plot, which shows that early all the reads begin after position 58, forming a “cliff”. (**c**) After AlkB demethylation, however, the read alignments lengthen and extend past the cliff, resulting in a more varied distribution of alignment start positions. The heat map shows a significant reduction in the number of aligned reads. (**d**) Many RNA modifications can also be mapped by polymerase-induced mutations in the resulting cDNA. This is illustrated with a striking T-to-C misreading in BCG tRNA Arg-ACG, which is consistent with the presence of inosine at position 34 of the anticodon on nearly all copies of the tRNA.

The presence of modified nucleotides appears to block RT, leading to a read interruption 1-2 nucleotides away from the modification site. Further validation of this idea comes from a BCG library preparation lacking AlkB demethylase. In the absence of AlkB, up to 90% of the reads mapping to 23 of 45 of tRNA species were truncated at positions 59-60, as illustrated in the heatmap of polymerase stops in **Figure 6b**. After AlkB treatment, most stops at positions 59-60 have disappeared and reads increased in length with a leftward shift toward the 5’ end of the tRNA (**Fig. 6c**). The alignment profile of Glu-CTC illustrates the effect of AlkB on the read distribution along the tRNA sequence. The sharp “cliff” at position 60 in the untreated sample (**Fig. 6b**) corresponds to a predicted m^1^A modification at position 58 (**Fig. 6b**). m^1^A is a known substrate for AlkB (**Fig. 1c**) and blocks RT.^15, 43, 44^ After AlkB treatment, the cliff disappears and the reads span the tRNA sequence (**Fig. 6c**). The presence of 5’ cliffs in the alignment plots for nearly all tRNA species in BCG (**Fig. S.1**) points to the potential for quantitative mapping of RNA modifications by AQRNA-seq.

Modification mapping is also illustrated by the presence of signature mutations resulting from RT.^44^ For example, inosine (I) is present at the wobble position of a single tRNA in mycobacteria^41^ and tends to pair with C during RT.^42, 45^ I is thus detected by a near stoichiometric T-to-C sequencing mutation, as illustrated in **Fig. 6d** for position 34 of BCG tRNA Arg-ACG. As reviewed extensively by Motorin and Helm, this kind of modification mapping could aid in the discovery of previously unannotated or unlocalized modifications in poorly characterized species. AQRNA-seq enables both modification mapping and mapping of cleavage sites while quantifying the landscape of the population of small RNA species in a single sample.

## Discussion

Here we have developed, validated, and applied the first RNA sequencing method that provides precise and accurate absolute quantification of all small RNA species in a single sample. The inability of current RNA-seq methods to provide accurate information about the quantities of individual RNA molecules in a sample mixture has constrained our understanding of RNA regulation and RNA biology. Following a systematic deconstruction of the RNA-seq NGS library prep workflow, we identified several steps critical to the quantitative precision and accuracy of RNA-seq results: (1) sequential ligation of the 5’ linker after reverse transcription, (2) linker structures and biochemical conditions that provide >90% ligation efficiency, (3) a non-essential but informative AlkB demethylation step to reduce interference from polymerase-blocking alkyl modifications, and (4) a data mining workflow that minimizes loss of read information to improve quantitative accuracy. Within AQRNA-seq, adapter ligation steps have been rigorously optimized to maximize RNA capture, with ligation of the 5’-adapter scheduled after reverse transcription to ensure that all cDNA species, including truncated cDNAs, are captured and represented in the final library mixture. A non-essential AlkB demethylation step is employed to minimize premature cDNA truncation and to provide information about the identities and locations of polymerase-blocking or mutating modifications. Beyond the linear relationship between read count and RNA copy number, the RNA-seq method provides information about modification occupancy, tertiary structure, and fragmentation.

While our RNA-seq method was applied first to tRNA analysis, it is broadly applicable to any form of RNA. The validation against the miRNA library illustrates this general applicability. The 963 miRNAs in the Miltenyi miRXplore Universal Reference possess varying lengths (16-28 nt) and varying sequences (including all 16 possible 3’- and 5’-termini), which provides a robust test of the quantitative accuracy of the RNA-seq method. With the best-performing commercial kits able to quantify only 40-50% of the miRNAs within a two-fold range of accuracy, the fact that we were able to quantify ∼75% of the miRNAs to within two-fold of expected abundance, without incidence of jackpot and a low incidence of dropouts, attests to the rigor of the RNA-seq method. The AQRNA-seq method is thus established to be the most accurate for application in miRNA profiling studies, with the recent miRNA method developed by Kim *et al*. lacking evidence of quantitative rigor.^14^ However, while random priming RNA-seq methods provide relatively accurate quantification of mRNAs,^46^ AQRNA-seq is also applicable to longer RNA species such as mRNA and rRNA as a means to map RNA modifications and cleavage sites, as illustrated in **Fig. S4**. The addition of a Covaris fragmentation step after ligation of the first linker allows longer RNA species to be reduced to an acceptable length for library generation. The introduction of this step allows for quantification of (1) all expressed copies of an RNA (all molecules with a 3’-end that maps to the end of the transcribed or fully mature sequence), (2) polymerase fall-off due to modifications, secondary structure, or polymerase detachment, and (3) fragmentation sites within the RNA molecules (3’-ends that map within the full-length sequence).

## Conclusion

Collectively, our results demonstrate a quantitatively accurate method for sequencing RNAs. We have rigorously validated the accuracy of the method to ensure ligation and amplification biases are minimized. In contrast to some currently available approaches, AQRNA-seq is generalizable to all classes of RNA – short noncoding RNAs such as microRNAs, highly-modified species such as tRNAs, and longer mRNAs, rRNAs, and other types of RNA.

## Acknowledgments

We thank Sidney Vermeulen for technical assistance during method development and members of the Bartel Laboratory (Whitehead Institute and MIT Department of Biology), especially Asia Stefano and Prof. David Bartel, for assistance with northern blots. We thank Dr. Pavel Ivanov (Harvard Medical School) for sharing synthetic RNA standards. This work was supported by grants from the US National Science Foundation (MCB-1412379 to V.C.L.), the National Institutes of Health (ES002109), and the National Research Foundation of Singapore through the Singapore-MIT Alliance for Research and Technology Antimicrobial Resistance IRG. J.F.H. was supported by the MIT Toxicology Training Grant T32-ES007020 to J.F.H.), D.Y. by a postdoctoral fellowship from A*STAR, Singapore, and S.M.H. by a postdoctoral fellowship from the Swiss National Science Foundation.

## Author Contributions

P.C.D., B.C., J.F.H. conceived the idea, designed the experiments, and wrote the first draft of the manuscript. P.C.D., B.C., D.Y. and J.F.H. developed the method and performed the sequencing experiments. J.F.H., B.C., D.M., and S.S.L. implemented and interpreted the computational analyses. D.Y. and J.F.H. performed mycobacterial culturing and RNA isolation. J.M.B. performed *E. coli* culturing and RNA isolation. S.M.H. optimized experimental conditions and characterized demethylation efficiencies by LC-MS. S.M.H. and J.F.H. performed northern blot analyses. M.S.D. contributed reagents. V.C.L. supervised *E. coli* experiments and contributed insights and analysis. All authors participated in the writing of the manuscript.

## Data Availability

The complete sets of data for the RNA sequencing studies reported in **Figures 3-6** have been deposited in the NCBI Sequence Read Archive (SRA) under the BioProject ID PRJNA579244. miRNA and standards data shown in **Figure 2** have been deposited in the Gene Expression Omnibus (GEO) under accession number GSE139936.

## Code Availability

The software used in the studies presented here is publicly available as follows. Blast version 2.6.0 (nucleotide BLAST) available at https://blast.ncbi.nlm.nih.gov/Blast.cgi?PAGE_TYPE=BlastDocs&DOC_TYPE=Download. Peakfit.m version 9.0 available at Tom O’Haver, MATLAB Central File Exchange - https://terpconnect.umd.edu/~toh/spectrum/. fgrep (Linux command) available at https://unix.stackexchange.com/questions/17949/what-is-the-difference-between-grep-egrep-and-fgrep. fastxtoolkit version 0.013 available at http://hannonlab.cshl.edu/fastx_toolkit/. Custom python scripts (.gitignore, blast_pair.py, cull.py) are available at GitHub https://github.com/dedonlab/aqrnaseq.

