## Supplementary material for "Sequencing-based quantitative mapping of the cellular small RNA landscape": Methods

**Bacterial strains, culturing conditions, growth assays, and RNA isolation**

All *E. coli* strains used were taken from the Keio collection (GenoBase; http://ecoli.aist-nara.ac.jp/)[1]. The genotype of each of the bacterial strains was validated before conducting the study. Strains were cultured in 10 ml LB broth (Fisher BioReagents BP1426-2) at 37 °C with constant shaking at 180 rpm until the cultures reached a final $OD_{600}$ of 0.6-0.7. Culture pellets were harvested by centrifugation at 13,500 rpm for 2 min and immediately used for tRNA isolation using the Purelink miRNA isolation kit (Thermofisher K15700) following the manufacturer's protocol. Briefly, cell pellets were resuspended in Trizol Reagent (Thermofisher 15596018) for lysis, followed by treatment with chloroform to separate the aqueous layer containing bulk tRNA. The aqueous layer was then subjected to a 2-column purification process where genomic DNA, larger RNA fragments (>200 bp), and excess salts, are removed. It is important to note that 5S rRNA cannot be separated from tRNA using this method. Three biological replicates were used in *E. coli* tRNA study.

*Mycobacterium bovis* Bacille Calmette-Guérin (BCG) str. Pasteur 1173P2 was grown in roller bottles with 7H9 broth or PBS (with 0.05% (v/v) tyloxapol, Sigma Aldrich T8761) at 2 rpm and 37 $^{o}$C. Exponentially growing cultures with an $OD_{600}$ of 0.8-1.0 were starved by washing pellets three times with PBS-tyloxapol. Starvation cultures were inoculated into PBS-tyloxapol at a starting $OD_{600}$ of 1.0. Samples were retrieved at 4, 10, and 20 d after starvation. At day 20, cultures were resuspended in 7H9 and resuscitated for 6 d prior to harvesting. At each time point, cultures were plated on 7H10 agar for CFU determination. Specific compositions of 7H9 media, PBS, and 7H10 agar are as follows. Middlebrook 7H9 (BD Difco 271310) was supplemented with 0.5% (w/v) albumin (Sigma Aldrich A3059), 0.2% (w/v) glucose (Sigma Aldrich G8270), 0.085% (w/v) NaCl (Sigma Aldrich S7653), 0.2% (v/v) glycerol and 0.05% (v/v) Tween 80 (Sigma Aldrich P8074) as nutrient replete media; PBS (137 mM NaCl , 2.7 mM KCl, 10 mM $Na_2HPO_4$, 2 mM $KH_2PO_4$) was supplemented with 0.05% v/v tyloxapol, a non-hydrolyzable detergent. 7H10 agar (BD Difco 262710) was supplemented with 0.5% (v/v) glycerol and 10% (v/v) oleic acid albumin dextrose catalase (OADC; BD BBL 212351).

For colony forming unit (CFU) determination, serial dilutions of BCG cultures at various nutrient starvation/resuscitation timepoints were plated on 7H10 agar and incubated at 37 $^{o}$C for 3-4 weeks. Colonies of the BCG timepoints were subsequently counted and respective CFUs determined.

For RNA isolation and purification from BCG, cells collected at nutrient starvation/resuscitation timepoints were lysed in the presence of TRI reagent (Sigma Aldrich T9424) with glass beads in a FastPrep FP120 bead-beater as previously described[2]. Small RNAs were subsequently isolated using the Purelink miRNA isolation kit (Invitrogen K157001). Small and large RNA fractions were profiled using the Agilent Bioanalyzer.

**Optimized AQRNA-seq protocol**

*Ligation of input RNA and DNA Linker 1.* Small RNAs (50 ng) were mixed with an 80-mer spike-in RNA oligo internal standard. The mixture was dephosphorylated in a 5 µl reaction containing 0.5 µl of reaction buffer (NEB T4 RNA ligase buffer) and 1 U of shrimp alkaline phosphatase (rSAP, NEB) at 37 °C for 30 min. The reaction was stopped by heat inactivation at 65 °C for 5 min followed by cooling on ice. Linker 1 ligation was performed by adding the following reagents directly to the dephosphorylation product: 1 µl Linker 1 (100 pmol/ul), 3 µl ATP (10 mM, NEB), 2.5 µl T4 RNA ligase buffer (NEB), 2 µl T4 RNA ligase 1 (30U/ul, NEB), 1.5 µl water and 15 µl PEG8000 (NEB). The ligation mixture was incubated at 25 °C for 2 h and 16 °C overnight. The ligation product was purified using the Zymo Oligo Clean & Concentrator kit (Zymo Research, D4060) according to the manufacturer's instructions. The sample was eluted in 20 µl water and kept on ice prior to Bioanalyzer analysis (Agilent, small RNA kit) or used directly in the demethylation step.

*Demethylation.* The ligated RNA sample was demethylated by AlkB (Arraystar, rtStar tRF&tiRNA Pretreatment Kit). A 2X reaction buffer was freshly prepared before the reaction and consisted of 150 µM 2-ketoglutarate, 4 mM L-ascorbic acid, 150 uM $(NH_4)_2Fe(SO_4)_2$, 100 µg/mL BSA (NEB, molecular biology grade, 10 mg/mL), and 100 mM HEPES (pH 8.0). The demethylation reaction was performed in a 100 µl volume consisting of 50 µl 2X reaction buffer, 20 µl Linker 1-ligated tRNA sample, 2 µl AlkB demethylase, and 1 µl RNase Inhibitor (NEB, murine, 40,000U/mL). The reaction was incubated at ambient temperature. After 2 h, the reaction was stopped via the addition of 50 µl water and 100 µl phenol:chloroform:isoamyl alcohol 25:24:1, pH 5.2. The mixture was mixed by inverting several times and centrifuged at 16,000xg for 10 min. The top layer was transferred to a new Eppendorf tube. Another 100 µl chloroform was added to the original mixture to remove any remaining phenol. After centrifugation, the top layer was removed and combined with the first extraction. The extracted sample was then purified using the Zymo Oligo Clean & Concentrator kit (Zymo Research, D4060) per the manufacturer's instructions. The sample was eluted in 17 µl water before proceeding to the next step (Linker 1 removal).

*Removal of excess DNA Linker 1.* In this step, the DNA oligo–adenylate intermediate was de-adenylated and subsequently digested, together with unused Linker 1, by exonuclease RecJ. The de-adenylation was performed in a 20 µl reaction containing 16 µl RNA sample from demethylation step, 2 µl NEB Buffer 2 (10X), and 2 µl 5'-deadenylase (NEB, 50U/uL). After incubation at 30 °C for 1 h, 2 µl RecJ (NEB; 30U/uL) was added. The mixture was incubated at 37 °C for 30 min followed by the addition of another 2 µl RecJ and further digestion for an additional 30 min. The reaction was stopped by heating at 65 °C for 20 min. The reaction mixture was purified using a DyEx spin column (Qiagen, 63204).

*Reverse transcription.* The RNA sample obtained from DyEx column purification is mixed with 1 µl RT-primer (2 pmol/ µl) and 1 µl dNTPs (10 mM each). The mixture was heated at 80 °C for 2 min and cooled immediately on ice. PrimeScript Buffer (6 µl; Clontech), 1 µl RNase Inhibitor (NEB), and 1 µl PrimeScript Reverse Transcriptase (Clontech, RR014A) were added. The reverse transcription reaction mixture was then incubated at 50 °C for 2 h after which the enzyme was inactivated at 70 °C for 15 min. The RNA strand was then hydrolyzed by adding 1 µl NaOH (5 M) followed by incubation at 90 °C for 3 min. The hydrolysis product was neutralized by adding 1 µl HCl (5M) and the reaction was cleaned up using the Zymo Oligo Clean & Concentrator kit (Zymo Research, D4060). The sample was eluted with 15 µl of water before vacuum concentration to 5 ul.

*cDNA ligation.* The purified cDNA was ligated to Linker 2 (**Table 2-4**) in a 20 µl reaction consisting of 5 µl cDNA sample, 1 µl Linker 2 (50 pmol/µl), 2 µl T4 DNA Ligase Buffer (NEB), 1 µl ATP (10 mM, NEB), 2 µl T4 DNA ligase (400U/uL, NEB), and 9 µl PEG8000 (NEB). The mixture was mixed and then incubated at 16 °C overnight for ligation. The ligated product was purified using the Zymo Oligo Clean & Concentrator kit (Zymo Research, D4060) per the manufacturer's instructions and eluted in 16 µl of water.

*Removal of excess DNA Linker 2.* After cDNA ligation, excess Linker 2 was removed by a two-step purification. First, adenylated linker intermediates were de-adenylated. Then, RecJ was used to digest the de-adenylated product. The de-adenylation was performed in a 20 μl reaction containing 16 μl RNA sample from the previous, cDNA ligation step, 2 μl NEB Buffer 2 (10X), and 2 μl 5'-deadenylase (NEB, 50U/ul). After incubation at 30 °C for 1 h, 2 μl RecJ (NEB; 30U/ul) was added. The reaction was incubated at 37 °C for 30 min. Subsequently, another 2 μl RecJ was added for further digestion for an additional 30 min. The reaction was stopped through heat inactivation at 65 °C for 20 min.

*PCR amplification and Illumina sequencing.* Purified cDNA from the previous step was amplified by PCR in a 100 μl mixture containing 24 μl cDNA template, 50 μl seqAMP DNA polymerase buffer (2X buffer, Clontech), 2 μl each of PCR primer F and R with unique sequencing barcodes (**Table 2-4**, 1 μM each), 2 μl Clontech seqAMP DNA polymerase (Clontech) and 20 μl water. The PCR reaction was performed according to the manufacturer's instructions with an annealing temperature of 58 °C and 13 reaction cycles. The PCR product was extracted and purified from an agarose gel using a standard gel purification kit (QIAquick Gel Extraction Kit, Qiagen). The gel-extracted samples were mixed together (multiplexing) and submitted for Illumina sequencing. In the studies described, sequencing was performed on the Illumina NEXTseq sequencer (BioMicroCenter, MIT) with custom primers F and R (**Table 1-1**).

**Table 1-1 DNA and RNA oligos required for the AQRNA-seq protocol.** Asterisk (*) = sites of phosphorothioate modification. N = randomized position (equal mixture of all 4 bases used during oligo synthesis). Unless otherwise indicated, oligos terminate in hydroxyl groups at both 5' and 3' ends.

| Name | Sequence (5'-3') |
|---|---|
| Spike-in internal RNA standard (80-mer) | ACCCACGGCAGAGUGUGUGUGGCCCACGCGAUUCGUGAAUAACAUA ACUAUGAGUAGGAUAAGGAAUGUCACCUAACAGC |
| Optional second spike-in internal RNA standard (40-mer) | NNNUAUCCUAAUCAUCUUAAUACUACAAUGGACCAUGNNN |
| Linker 1 | /5Phos/-NNCAC TCG GGC ACC AAG GAddC-3' |
| Linker 2 | /5Phos/TG AAG AGC CTA GTC GCT GTT CAN NNN NNC TGC CCA TAG AG/3SpC3/ |
| RT-primer | G*T*C*C*T*T*GGTGCCCGAGTG |
| PCR primer F-1 | AATGATACGGCGACCACCGAGATCTACACAGAGAGACACTCTTTCCCT ACACGACGCTCTTCCGATCTTGAACAGCGACTAGGCTCTTCA |
| PCR primer F-2 | AATGATACGGCGACCACCGAGATCTACACGTGTGTACACTCTTTCCCT ACACGACGCTCTTCCGATCTTGAACAGCGACTAGGCTCTTCA |
| PCR primer F-3 | AATGATACGGCGACCACCGAGATCTACACTCTCTCACACTCTTTCCCT ACACGACGCTCTTCCGATCTTGAACAGCGACTAGGCTCTTCA |
| PCR primer F-4 | AATGATACGGCGACCACCGAGATCTACACCACACAACACTCTTTCCCT ACACGACGCTCTTCCGATCTTGAACAGCGACTAGGCTCTTCA |
| PCR primer F-5 | AATGATACGGCGACCACCGAGATCTACACCAAGGTACACTCTTTCCCT ACACGACGCTCTTCCGATCTTGAACAGCGACTAGGCTCTTCA |
| PCR primer F-6 | AATGATACGGCGACCACCGAGATCTACACAGGTTCACACTCTTTCCCT ACACGACGCTCTTCCGATCTTGAACAGCGACTAGGCTCTTCA |
| PCR primer R-1 | CAAGCAGAAGACGGCATACGAGATCCTGAGCGGTCTCGGCATTCCTG CTGAACCGCTCTTCCGATCTGTCCTTGGTGCCCGAGTG |
| PCR primer R-2 | CAAGCAGAAGACGGCATACGAGATTGCTGTCGGTCTCGGCATTCCTG CTGAACCGCTCTTCCGATCTGTCCTTGGTGCCCGAGTG |
| PCR primer R-3 | CAAGCAGAAGACGGCATACGAGATCATCACCGGTCTCGGCATTCCTG CTGAACCGCTCTTCCGATCTGTCCTTGGTGCCCGAGTG |
| PCR primer R-4 | CAAGCAGAAGACGGCATACGAGATTTCAGACGGTCTCGGCATTCCTG CTGAACCGCTCTTCCGATCTGTCCTTGGTGCCCGAGTG |
| PCR primer R-5 | CAAGCAGAAGACGGCATACGAGATACGGTGCGGTCTCGGCATTCCTG CTGAACCGCTCTTCCGATCTGTCCTTGGTGCCCGAGTG |
| PCR primer R-6 | CAAGCAGAAGACGGCATACGAGATGTAACACGGTCTCGGCATTCCTG CTGAACCGCTCTTCCGATCTGTCCTTGGTGCCCGAGTG |
| Custom primer F | GCTCTTCCGATCT TGAACAGCGACTAGGCTCTTCA |
| Custom primer R | TGAACCGCTCTTCCGATCT GTCCTTGGTGCCCGAGTG |

**Optimization of AlkB demethylation conditions**

*LC-MS/MS analysis of methylated ribonucleosides.* Ribonucleosides were resolved with a Phenomenex Synergi Fusion-reverse phase column (100 x 2 mm, 2.5 µm particle size, 100 Å pore size) eluted with the following gradient of acetonitrile in 5 mM ammonium acetate (pH 5.3) at a flow rate of 0.35 ml/min and 35 °C: 0-1 min, 0%; 1-10 min, 0-10%, 10-14 min, 10-40%, 14-15 min, 40-80%. The HPLC column was coupled to an Agilent 6430 triple quadrupole LC/MS spectrometer with an electrospray ionization source where it was operated in positive ion mode with the following parameters for voltages and source gas: gas temperature, 350 °C; gas flow, 10 l/min; nebulizer, 45 psi; and capillary voltage, 3500 V. The first and third quadrupoles (Q1 and Q3) were fixed to unit resolution and the modifications were quantified by pre-determined molecular transitions. The dwell time for each ribonucleoside was 500 ms. The retention time, *m/z* of the transmitted parent ion, *m/z* of the monitored product ion, fragmentor voltage, and collision energy of each modified nucleoside are as follows: $m^1A$, 3.6 min, *m/z* 282 → 150, 100 V, 16 V; $m^1G$, 6.1 min, *m/z* 298 → 166, 90 V, 10 V; $m^1I$, 5.9 min, *m/z* 283 → 151, 80 V, 10 V; $m^{22}G$, 7.8 min, *m/z* 312 → 180, 100 V, 8 V. Modified ribonucleosides were identified using commercially available nucleoside standards. Three independent tRNA replicates were used to test AlkB demethylation efficiencies in this study.

*LC-MS/MS data analysis.* Quantitative comparisons between control and demethylase-treated samples from different buffers were made possible by correcting for variation in tRNA quantities by dividing raw peak are for each ribonucleoside by the ultraviolet absorbance peak areas for the four canonical ribonucleosides. The demethylation efficiencies were calculated by dividing the peak area of the demethylase-treated sample by the peak area of the control sample for each modification.

**Optimization of linker ligation conditions and determination of linker removal**

*Linker 1 ligation studies.* Tests of Linker 1 ligation efficiencies were performed by combining dephosphorylated tRNA with ATP, T4 RNA ligase buffer, T4 RNA ligase 1, PEG8000, and varying amounts of Linker 1. The reaction was allowed to proceed under conditions consistent with the manufacturer's recommendations and the resulting products were analyzed using the Bioanalyzer small RNA chip. Electropherogram peaks corresponding to ligated and unligated tRNA were fitted and integrated using Peakfit.m (version 9.0; Tom O'Haver, MATLAB Central File Exchange - https://terpconnect.umd.edu/~toh/spectrum/), a Matlab-based peak fitting program that uses an unconstrained non-linear optimization algorithm to decompose a complex peak signal into its fundamental underlying component parts. The fraction of ligated tRNA was calculated by dividing the summed peak area corresponding to the ligated tRNA to the total peak area. Three independent replicates were used for linker 1 ligation efficiency testing.

*Linker 2 ligation studies.* To test the efficiency of the Linker 2 ligation reaction, we used single-stranded DNA oligos with phosphorothioate modification at the 5' end to simulate the reverse transcription output. The sequences for the oligos used here are listed in **Table 2-4**. To determine the optimal linker:oligo ratio, we combined the 80-nucleotide oligo with ATP, T4 DNA ligase buffer, T4 DNA ligase, PEG8000, and varying amounts of Linker 2. To determine whether oligo length altered ligation efficiencies, we used a constant linker:oligo ratio and varied the length of the oligo tested. The reactions were allowed to proceed under conditions consistent with the manufacturer's recommendations and the resulting products were analyzed using the Bioanalyzer small RNA chip. Similar to Linker 1 ligation studies, peaks were deconvoluted and integrated using Peakfit.m. The fraction of ligated oligo was used as a metric for ligation efficiency and used to determine final, optimized ligation conditions. Three independent replicates were used for cDNA ligation efficiency testing.

**Table 2-4 DNA oligo sequences for Linker 2 ligation studies.** Asterisk (*) = sites of phosphorothioate modification. N = randomized position (equal mixture of all 4 bases used during oligo synthesis). Unless otherwise indicated, oligos terminate in hydroxyl groups at both 5' and 3' ends.

| Name | Sequence (5'-3') |
|---|---|
| cDNA 50-mer | G*G*T*A*G*C*ACAAAAGTATTACCATGGTCCTAGAAGTTCGGCACAGTTANNN |
| cDNA 60-mer | A*C*T*G*T*T*GTTTGGTAGCACAAAAGTATTACCATGGTCCTAGAAGTTCGGCACAGTTANNN |
| cDNA 80-mer | A*C*G*C*T*G*CCACGTGTTCATTAACTGTTGTTTGGTAGCACAAAAGTATTACCATGGTCCTAGAAGTTCGGCACAGTTANNN |

*Linker removal studies.* The linker removal steps were performed on samples from the Linker 1 and 2 ligation studies. The extent of linker removal by deadenylase and RecJ was determined from Bioanalyzer electropherograms using peak fitting and integration. Three independent replicates were used to test linker 1 and 2 removal efficiencies in this study.

### Library construction for control AQRNA-seq samples

*Standard RNA oligos.* Five synthetic RNA oligos of different lengths and sequences were used to study the effect of oligo concentration and length on sequencing response. The oligos, whose sequences are listed in **Table 2-5**, were diluted from stocks to varying final concentrations and mixed together into standard mix samples A-E according to the scheme presented in **Table 2-6**. Samples A through E were prepared in triplicate and used as input RNA samples for AQRNA-sequencing.

**Table 2-5 RNA oligo sequences for linearity studies.** N = randomized position (equal mixture of all 4 bases used during oligo synthesis). Unless otherwise indicated, oligos terminate in hydroxyl groups at both 5' and 3' ends.

| Name | Sequence (5'-3') |
|---|---|
| 25-mer std | NNCAUUGUUAAGCGAACGGCUGCNN |
| 40-mer std | NNNUAUCCUAAUCAUCUUAAUACUACAAUGGACCAUGNNN |
| 60-mer std | NNNCGACCCAAGUUGUCUGAUUAAGCAUAUUAAGAGCCGAGGAUCGUAUCG CCACCCNNN |
| 80-mer std 1 | UCCUGAUAAUUGCAGAGAAGUGACUUCGUGGGUGUCACCAGUAAUAAGGCA UCGCCAGCUAACAAUCGGUGGGAACGUCA |
| 80-mer std 2 | UGUCUUUCACGGAGACUGCGGGUAGCUCUGACAAGCUCCUAUGGUGAUACC ACCACGCUCGUAAUGAGAUGGGAUAGUCG |

**Table 2-6 Standard mixes of RNA oligos.** Target abundances (in femtomoles) of each standard within the five standard mix samples A, B, C, D, and E.

| | A | B | C | D | E |
|---|---|---|---|---|---|
| 25-mer std | 40 | 80 | 120 | 160 | 200 |
| 40-mer std | 80 | 120 | 160 | 200 | 40 |
| 60-mer std | 120 | 160 | 200 | 40 | 80 |
| 80-mer std 1 | 160 | 200 | 40 | 80 | 120 |
| 80-mer std 2 | 200 | 40 | 80 | 120 | 160 |

*MicroRNA mixture*. The miRXplore Universal Reference (Miltenyi Biotec) consists of 963 synthetic unmodified, HPLC-purified RNA oligos. The sample was reconstituted according to manufacturer's instructions and aliquoted. Three aliquots were used as separate RNA inputs for AQRNA-seq library construction.

### AQRNA-seq data processing

Forward and reverse reads were trimmed of their respective adapter sequences using fastxtoolkit (version 0.013). A minimum adapter alignment length of 10 bp was required, and unknown (N) nucleotides were kept. Sequences were blasted against a reference library using blast (version 2.6.0) with the parameters: blast -perc identity 90 -word_size 9 -dust no -soft_masking false. For sequencing libraries prepared from BCG tRNA, a reference library was created based on the 48 entries in the genomic tRNA database for *Myocbacterium bovis* BCG str. Pasteur_1173P2.[3] (http://gtrnadb.ucsc.edu/GtRNAdb2/genomes/bacteria/Myco_bovi_BCG_Pasteur_1173P2_BCG_Pasteur_1173P2/Myco_bovi_BCG_Pasteur_1173P2_BCG_Pasteur_1173P2-gene-list.html). Sequences corresponding to duplicate tRNA genes (e.g., tRNA-Ala-TGC-1-2 and tRNA-Ile-GAT-1-2) and tRNA pseudogenes (tRNA-Ser-CGA-2-1) were removed to eliminate redundant entries and reduce the incidence of ambiguous or false positive matches. The terminal (3') CCA sequence was added to tRNA sequences where it is not genomically encoded. The sequence for the 80-nt RNA oligo internal standard was added to the reference library. For the control samples, the sequences of the 5 synthetic RNA oligos was used to create a reference library, along with the 80-nt RNA oligo internal standard.

For each tRNA and control sample, forward and reverse reads were merged by integrating their start and end positions to generate new start and end positions that reflect their combined coverage. Multiple alignments were reduced by ranking all the alignments for a given read by their e-value and retaining only the alignment with the lowest e-value. Forward and reverse reads were required to match the same target. Paired reads that did not match the same target were stored in a separate file and not analyzed. These manipulations were carried out with the python script cull.py. Uniquely-mapped reads were then tabulated and counted. For the microRNA samples, the set of 963 microRNA sequences contained within the miRXplore Universal Reference product was combined with the sequence for the 80-nt RNA internal standard oligo to generate the reference library. In that analysis, the number of exact sequencing reads that matched to the reference microRNA sequence in each trimmed sequencing file was determined with fgrep: numberReadsPerFile=$(fgrep $miRNA_sequence $trimmed_sequencing_file | wc -l). The read counts of the miRNAs were normalized to the summed counts for all detected miRNAs to obtain a “normalized read count”. The summed counts of all detected miRNAs were also divided by 963, the total number of detected miRNAs, to obtain the “expected read count” assuming all species were equimolar. The read ratio was calculated by dividing the normalized read count by the expected read count.

**Data availability**

All custom scripts have been made available at https://github.com/dedonlab/aqrnaseq. Additional modified scripts can be accessed upon request. All sequencing data that support the findings of this study have been variously deposited in the National Center for Biotechnology Information Sequence Read Archive (BioProject accession number PRJNA579244; BCG starvation data files) and the Gene Expression Omnibus (GEO accession number GSE139936; control and microRNA data files). All other relevant data are available from the corresponding author upon request.
