## Supplementary Information for "Sequencing-based quantitative mapping of the cellular small RNA landscape"

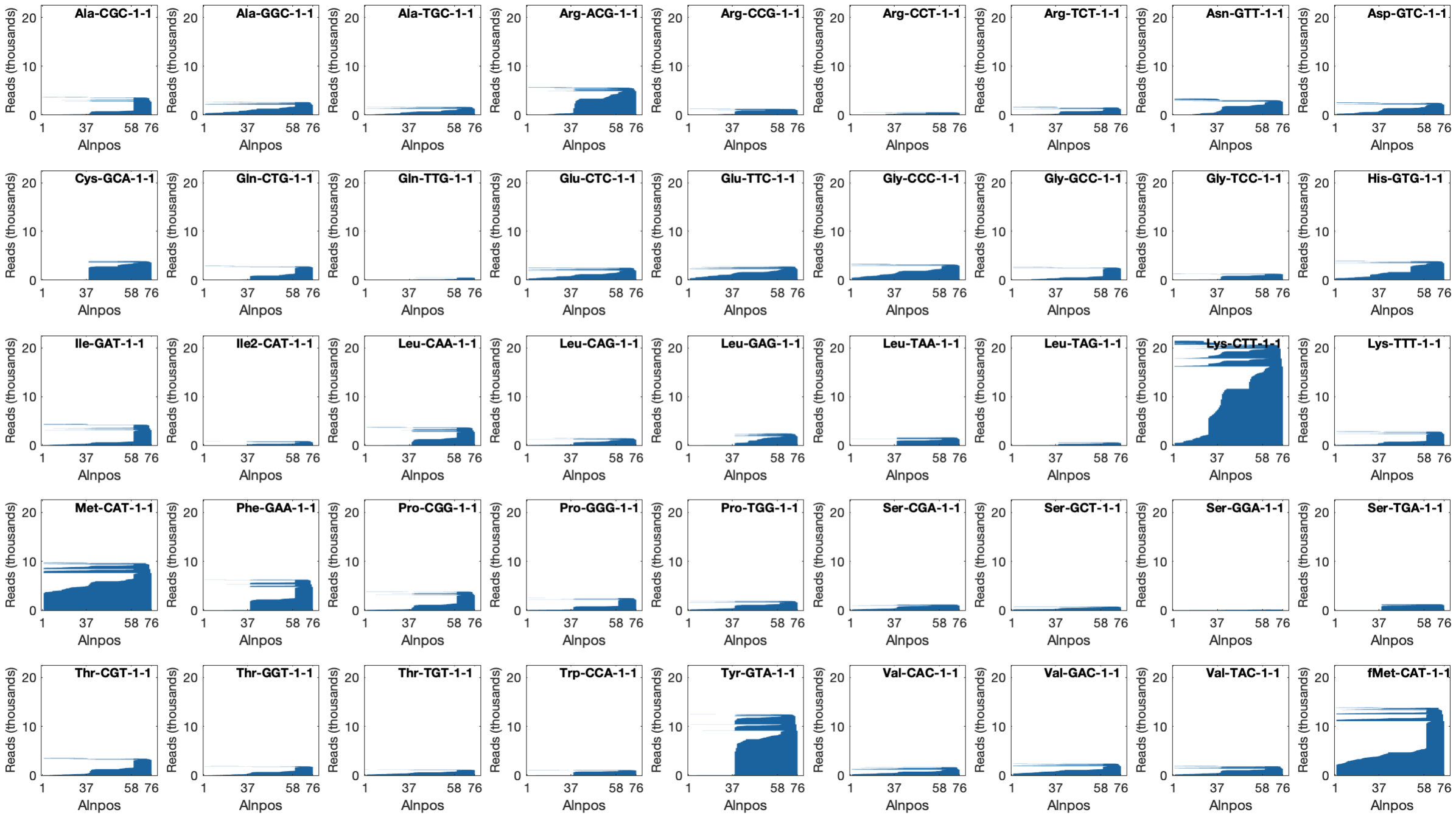

**Figure S1.** Complete set of alignment plots for the set of 45 unique, annotated tRNA genes from *M. bovis* BCG during log growth in rich medium. The y-axes of the plots are scaled to the highest expressed tRNA gene, tRNA Lys-CTT.

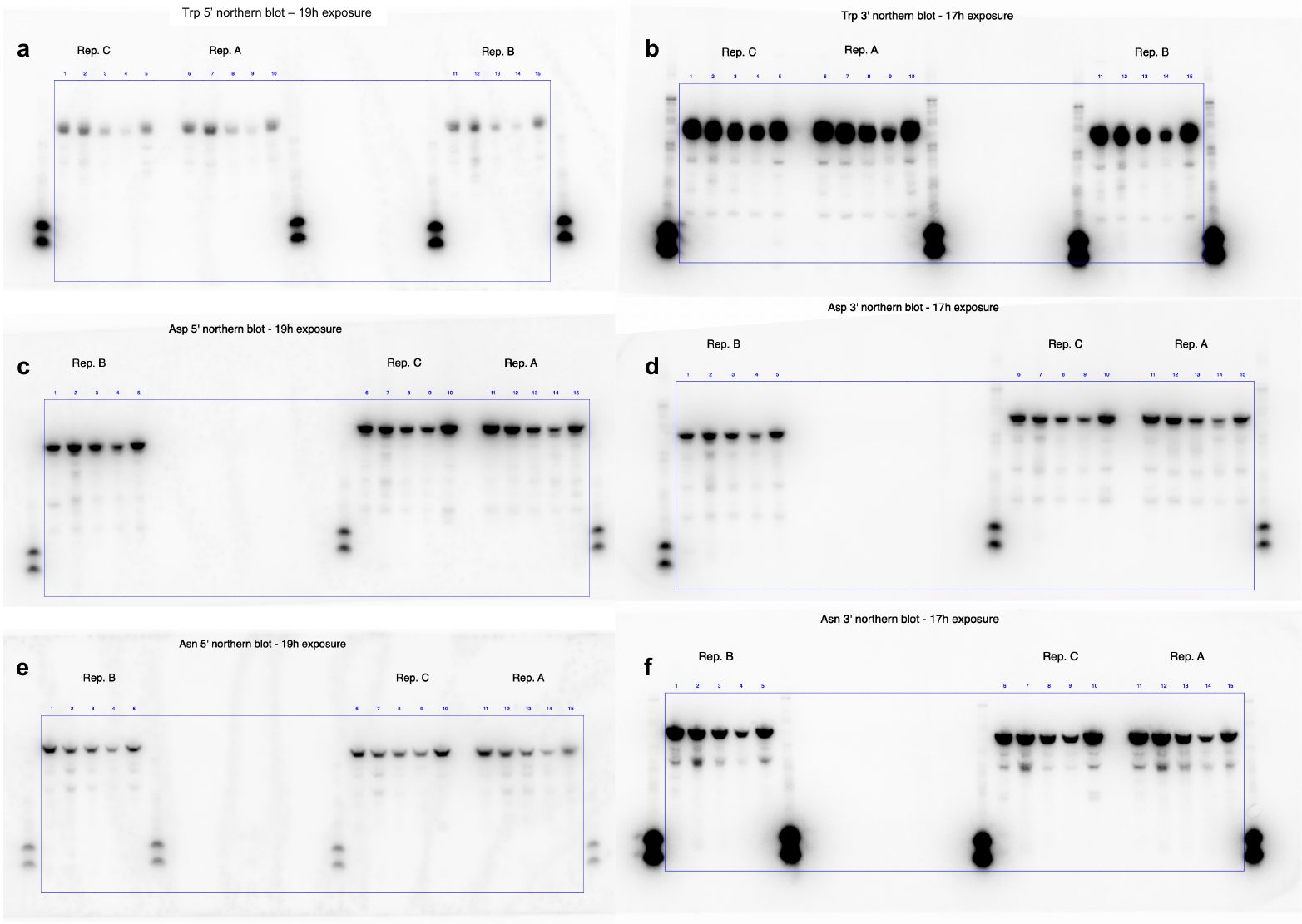

**Figure S2.** Northern blots performed using 5’ and 3’ probes against tRNA Trp-CCA (a, b), tRNA Asp-GUC (c, d), and tRNA Asn-GUU (e, f). Samples were loaded in the order of S0, S4, S10, S20 and R6 into lanes 1-5, 6-10 and 11-15. Each lane set corresponds to tRNA isolated from three biological replicates.

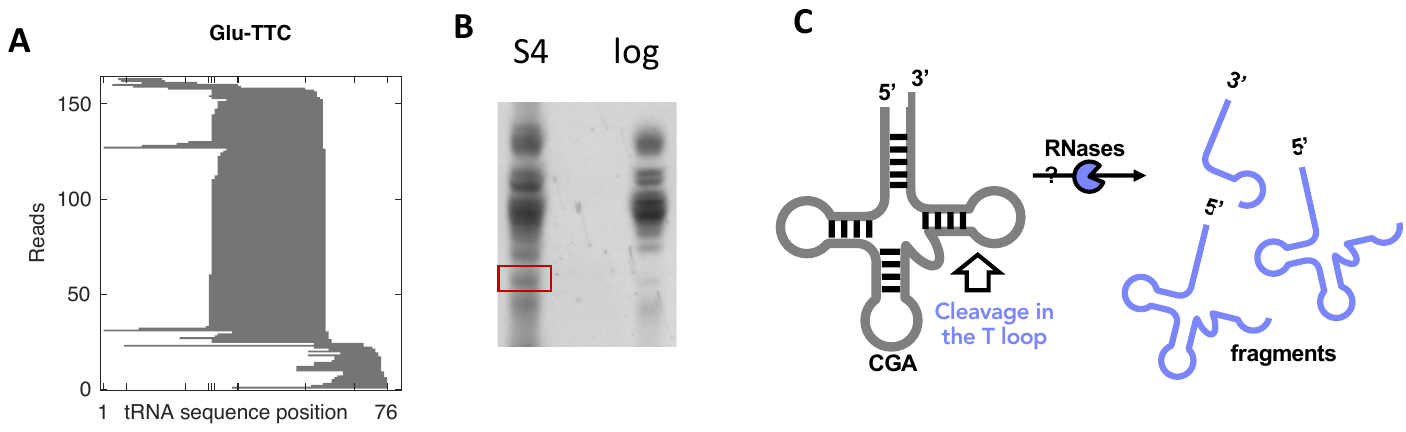

**Figure S3.** Small RNA was isolated from *M. bovis* BCG in log growth in rich medium (log) and on day 4 of starvation in PBS (S4). (**A**) Stack plot for AQRNA-seq analysis of small RNAs from S4. The corresponding stack plot for log small RNAs is shown in **Supplementary Figure S1**. (**B**) RNA from log and S4 cells was resolved on non-denaturing PAGE. The red box indicates location of a gel band that was subsequently excised and RNA purified for AQ-RNA-seq analysis. (**C**) Cleavage or degradation of tRNA Glu-TTC at starvation day S4 produces a fragment with the 3’ end located in the T loop of the tRNA structure.

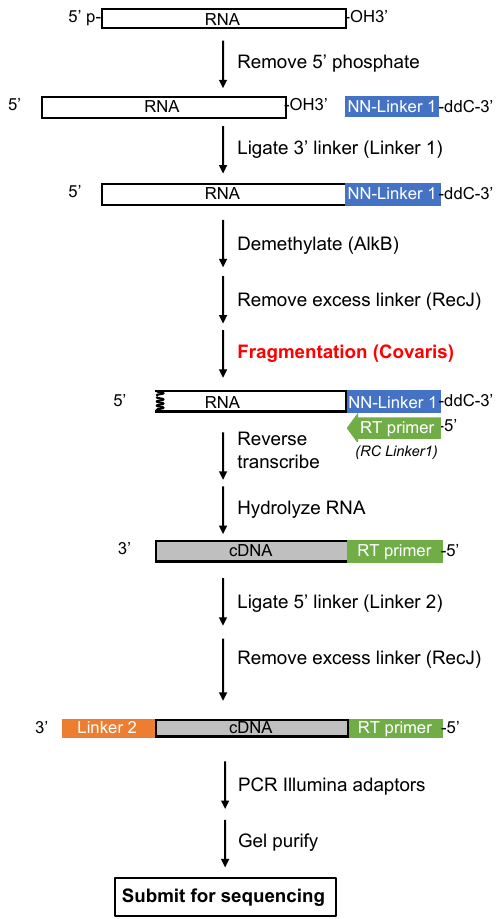

**Figure S4.** Schematic demonstrating a modified workflow for RNA species longer than 200 nt. The introduction of a Covaris fragmentation step after linker 1 ligation is highlighted.
